# Production of biologically active human basic Fibroblast Growth Factor (hFGFb) using *Nicotiana tabacum* transplastomic plants

**DOI:** 10.1101/2024.01.09.574869

**Authors:** Carolina Müller, Nicolás Budnik, Federico Gabriel Mirkin, Catalina Francisca Vater, Fernando Félix Bravo-Almonacid, Carolina Perez-Castro, Sonia Alejandra Wirth, María Eugenia Segretin

## Abstract

The use of plants as biofactories presents as an attractive technology with the potential to efficiently produce high-value human recombinant proteins in a cost-effective manner. Plastid genome transformation stands out for its possibility to accumulate recombinant proteins at elevated levels. Of particular interest are recombinant growth factors, given their applications in animal cell culture and regenerative medicine. In this study we produced recombinant human Fibroblast Growth Factor (rhFGFb), a crucial protein required for animal cell culture, in tobacco chloroplasts. We successfully generated two independent transplastomic lines that are homoplasmic and accumulate rhFGFb in their leaves. Furthermore, the produced rhFGFb demonstrated its biological activity by inducing proliferation in HEK293T cell lines. These results collectively underscore plastid genome transformation as a promising plant-based bioreactor for rhFGFb production.

**Main conclusion:** We generated transplastomic tobacco lines that stably express a human Basic Fibroblast Growth Factor (hFGFb) in their chloroplasts stroma and purified a biologically active recombinant hFGFb.

## Introduction

The use of plant-based bioreactors stands as an attractive method for the production of recombinant proteins for multiple applications (Tschofen et al. 2016). Plant molecular farming offers the advantage of cost-effective and large-scale production of heterologous proteins leveraging the abundance of plant biomass and the feasibility of utilizing established agricultural procedures for scaling-up (Schillberg et al. 2019; Shanmugaraj et al. 2020). Furthermore, the risk of contamination by endotoxins or animal pathogens is significantly diminished when compared to conventional expression platforms such as *Escherichia coli* or animal cell bioreactors (Buyel 2019; Clark and Maselko 2020). At present, there are successful developments of plant-produced recombinant proteins that have reached the market. Noteworthy examples include avidin (Merck, Germany), as well as the enzymes glucocerebrosidase and pegunigalsidase alfa (Protalix Biopharmaceuticals, Israel), among others, as comprehensively reviewed (Eidenberger et al. 2023).

A particularly promising plant-based expression system is based on the stable transformation of the plastid genome. In this system, transgene integration occurs in a specific site within the plastid genome through homologous recombination, thereby avoiding possible positional effects commonly associated with nuclear transformation (Maliga 2004). Given the polyploidy of chloroplasts, the transgene can exist in multiple copies per cell (Daniell 2006). These characteristics, coupled with the absence of reported gene silencing in chloroplasts, yield the potential for a high level of recombinant protein production. By means of example are the expression of the phage lytic protein (Oey et al. 2009) and the VP1-GUS protein (Lentz et al. 2010)—an epitope for the foot and mouth disease virus fused to β-glucuronidase—which represented 70% and 51% of total soluble proteins (TSP), respectively. An additional advantage of this expression system is the easy genetic manipulation and scaling-up when using *N. tabacum* as a host plant (Shanmugaraj et al. 2020).

Increasing interest has been directed towards recombinant growth factors due to their diverse applications, which include regenerative medicine, stem cell research and the emerging field of cell-culture meat production (Venkatesan et al. 2022). In particular, the human basic Fibroblast Growth Factor (hFGFb) is a key cytokine which regulates cell proliferation and pluripotency and is required for the maintenance of mammalian cell cultures (Levenstein et al. 2006; Lee et al. 2012; Mossahebi-Mohammadi et al. 2020).

The utilization of rhFGFb entails substantial costs for cell culture applications, encouraging the development of expression platforms that enable a cost-effective production. Within the field of molecular farming, different plant-based bioreactors were evaluated for the production of a biologically active rhFGFb, including transgenic cell lines from *Oryza sativa* and transgenic seeds from soybean *Glycine max* (Ding et al. 2006), *O. sativa* (An et al. 2013) and *N. benthamiana* (Yang et al. 2018). The production of rhFGFb in these systems was hindered either by relatively low expression levels or purifications yields of the recombinant protein.

Considering the remarkable potential of plastid transformation for achieving high levels of recombinant protein accumulation and the fact that hFGFb is a non-glycosylated protein, we chose this system for expressing rhFGFb in tobacco plants. We generated transplastomic plants that exhibited stable accumulation of a biologically active rhFGFb within the chloroplasts. Our findings demonstrate the promising potential of plastid genome transformation as an effective platform for producing recombinant rhFGFb, holding substantial implications for application in medicine, stem cell research and cellular agriculture.

## Results

### Transformation of tobacco plastid genome for rhFGFb expression results in homoplasmic plants

For the genetic transformation of tobacco plastid genome, we subcloned a synthetic 6xHIS tagged version of rhFGFb in the previously validated chloroplast transforming vector pBSWUTR (Wirth et al. 2004; Segretin et al. 2012) to obtain plasmid pUTR-rhFGFb (Fig.1a). The rhFGFb coding sequence was placed downstream of the promoter and 5’ untranslated region (5’UTR) of *psbA* gene (Eibl et al. 1999; Fernández-San Millán et al. 2003). The vector contains a selectable marker gene (*aadA*) that confers spectinomycin resistance to transplastomic shoots. This vector allows the integration of transgenes into the *rrn* operon, in the intergenic region located between the ribosomal *16s* and the *trn*I genes (Fig.1b). After transformation of *N. tabacum* leaves by particle bombardment with pUTR-rhFGFb, we selected nine spectinomycin resistant shoots and analyzed the transgene integration by PCR. The positive plants (Suppl. Fig. S1) were subjected to additional regeneration rounds in spectinomycin-containing medium to obtain homoplasmy. The transplastomic shoots from the third regeneration round were transferred to soil to obtain seeds.

**Fig 1.**
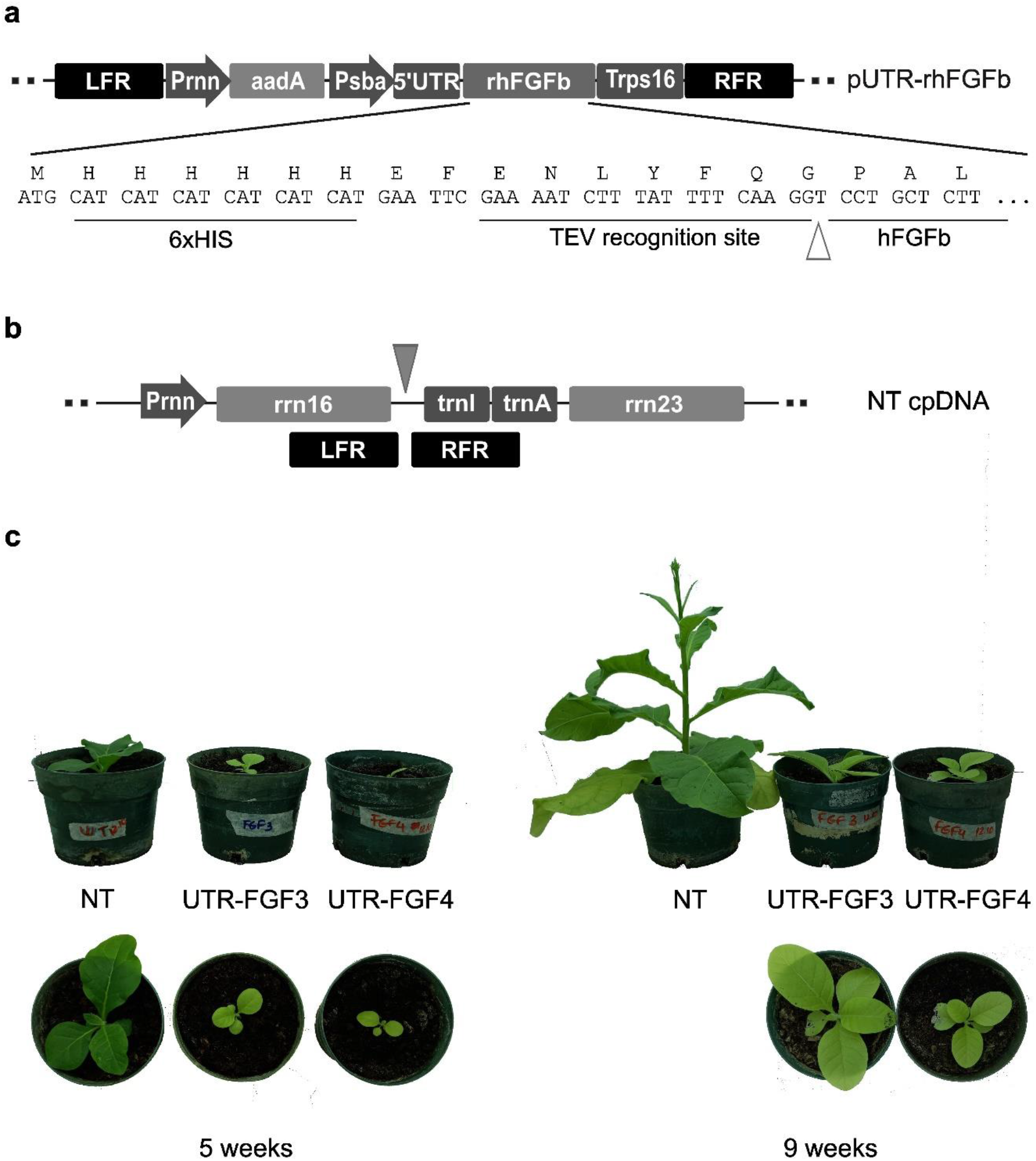
Generation of transplastomic plants for production of rhFGFb. **a** pUTR-rhFGFb vector containing the recombinogenic sequences flanking the selective gene (*aadA*) and the transgene (*hFGFb*). Left flanking region (LFR): 3’ region of the 16S ribosomal RNA sequence (*rrn16)*, Right flanking region (RFR): *trnI* and 5’ region of *trnA genes*. The rhFGFb transgene includes an open reading frame encoding a N-terminal Poly HIS-tag (6xHIS) followed by the recognition site of Tobacco Etch Virus protease (TEV) and the coding region of human FGFb. **b** Schematic representation of the integration site in the plastome. Transgenes are integrated in the intergenic region between *rrn16* and *trnI* (grey arrow). **c** Phenotype of transplastomic plants. Pictures of representative plants for each transplastomic line and the NT control taken after 6 and 9 weeks of germination.

Among 5 lines cultivated under greenhouse conditions, only two, designated UTR-FGF3 and UTR-FGF4, produced fertile seeds. These lines exhibited a slower growth rate and delayed flowering compared to non-transformed (NT) plants, while displaying a similar germination rate (Table 1). We recorded both plant size and number of fully expanded leaves per plant over time and find a developmental delay of approximately four weeks between UTR-FGF and NT plants. Furthermore, transplastomic plants exhibited a remarkable chlorotic phenotype in their leaves at all stages of development (Fig. 1c). This phenotype is associated with a reduction in chlorophyll levels, as measured in chlorophyll fluorescence units (SPAD) (Table 1).

**Table 1:** Phenotypic characterization of transplastomic lines UTR-FGF3 and UTR-FGF4. Plants were grown in soil and development and flowering was registered (n=4). Germination rate was measured *in vitro*. Chlorophyll fluorescence was measured from first and second fully expanded leaves. Values are reported as mean ± standard deviation of 3 independent experiments (germination rate) and at least 11 independent plants (chlorophyll fluorescence). Different letters indicate significant differences analyzed by Tuckey test (P<0.05).

| Line | Germination rate (%) | Number of leaves (6 week old plants) | Flowering time (weeks post germination) | Chlorophyll fluorescence (SPAD) |
| --- | --- | --- | --- | --- |
| NT | 92,7 ± 6,9 (a) | 5 | 12 | 27,26 ± 4,54 (a) |
| UTR-FGF3 | 93,4 ± 5,7 (a) | 4 | 20 | 10,02 ± 2,02 (b) |
| UTR-FGF4 | 93,9 ± 3,9 (a) | 4 | 20 | 10,40 ± 2,39 (b) |

We performed a Southern blot assay to confirm transgene integration and to determine the homoplasmic state (absence of non-transformed plastome copies). After extracting total DNA from leaves, we digested it with the restriction enzyme NcoI, which has two recognition sites flanking the integration site of the transgene. The used probe allows the distinction between the transformed (8.3 kpb) and non-transformed plastome (6.4 kpb). Both analyzed UTR-FGF lines show a single 8.3 kbp band indicating their homoplasmy (Fig. 2a).

**Fig 2.**
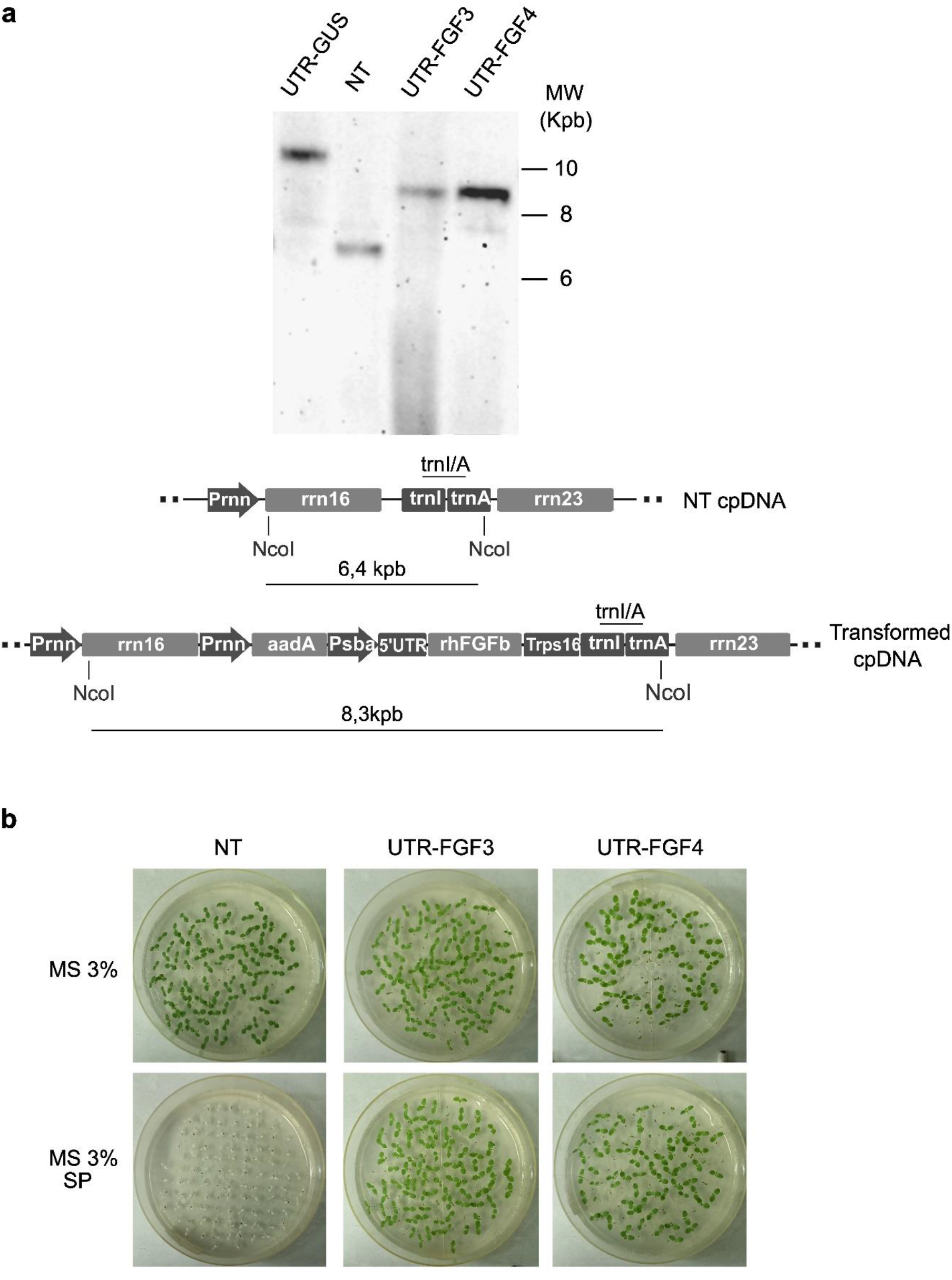
Analysis of homoplasmy and transgene inheritance. **a** Southern blot analysis (left) and schematic representation of transformed and non-transformed cpDNA (right) showing the restriction enzyme site and the trnI/A probe used for DNA hybridization. **UTR-GUS:** non-related transplastomic plant, **NT:** non-transfomed *N. tabacum*, **UTR-FGF:** transplastomic lines. **b** Germination assay: 100 seeds from each line were germinated in MS medium (MS) with or without spectinomycin (SP). Pictures were taken after 7 days.

Homoplasmy was further analyzed by evaluating the maternal inheritance and absence of transgene segregation. Seeds from transplastomic and NT plants were germinated in the presence of spectinomycin. As depicted in Fig. 2b, all transplastomic seedlings exhibited a green phenotype, indicating the absence of non-transformed chloroplast genome copies in the parental plants. These results reinforce the conclusion that the UTR-FGF lines are homoplasmic.

### rhFGFb accumulates in UTR-FGF lines and can be purified from leaves by affinity chromatography

To analyze the expression and accumulation of rhFGFb in leaves from transplastomic plants, we carried out a Western blot analysis. We observed a band of the expected size (18kDa) in the protein extracts obtained from the two transplastomic lines (Fig. 3a) confirming the expression of recombinant hFGFb. Furthermore, we analyzed the rhFGFb levels in leaves from different ages and determined that the developmental stage of the plant did not exert a significant impact on rhFGFb accumulation levels (Fig. 3b).

**Fig 3.**
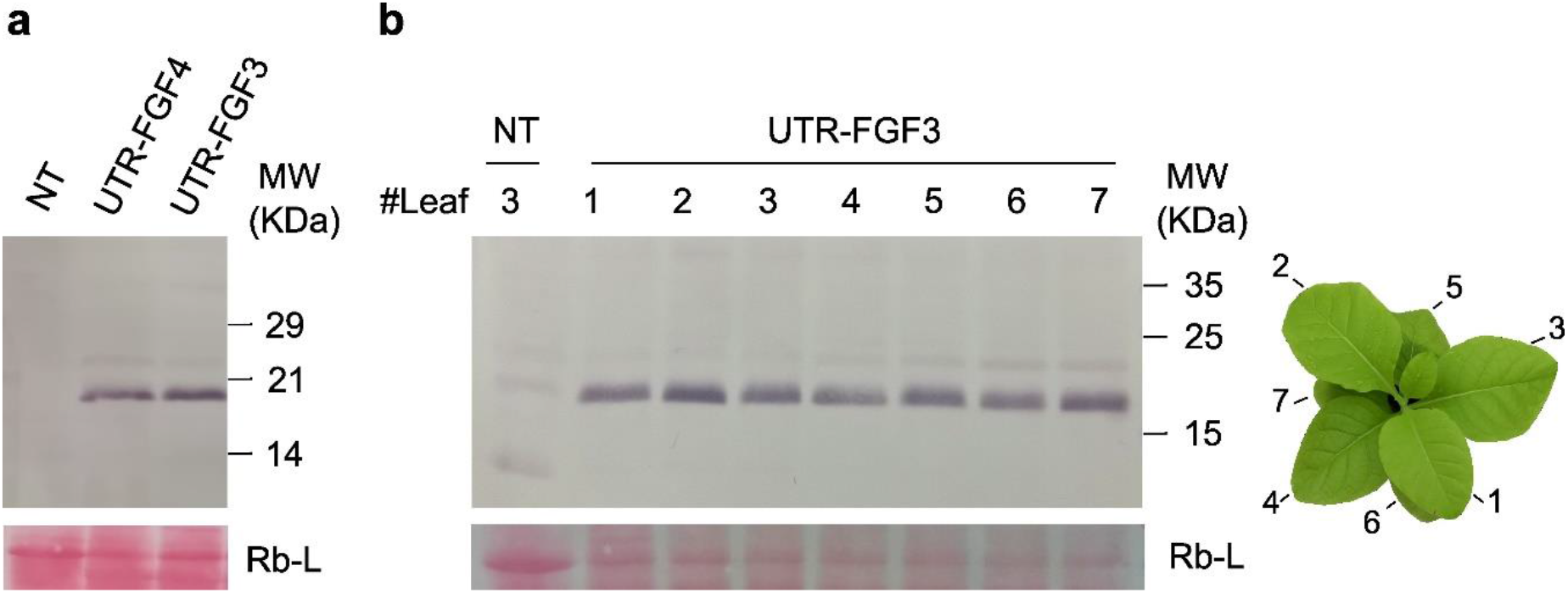
Analysis of protein accumulation in UTR-FGF transplastomic lines. Western blot analysis using anti-HIS antibody of **a**) rhFGFb accumulation in UTR-FGF lines and **b**) rhFGFb accumulation in leaves from different developmental stages. The different leaves used for protein extraction are marked with numbers starting from the first fully unfolded leaf (**b**, right). The RuBisCo large subunit (Rb-L) was used as a loading control (ponceau red dyed)

The recombinant protein was purified from fresh UTR-FGF3 leaves by Ni-NTA affinity chromatography (see Methods and Fig. 4a). Throughout the purification process, we detected a distinct band in the lanes corresponding to elution fractions 3 and 4, displaying a molecular weight consistent with the expected size of his-tagged rhFGFb. We confirmed its identity by Western blot analysis (Fig. 4c, lanes E3 and E4). The E3 fraction was filter-sterilized, and the buffer composition was adjusted to be equivalent to that of the commercial rhFGFb, for subsequent activity assays.

**Fig 4.**
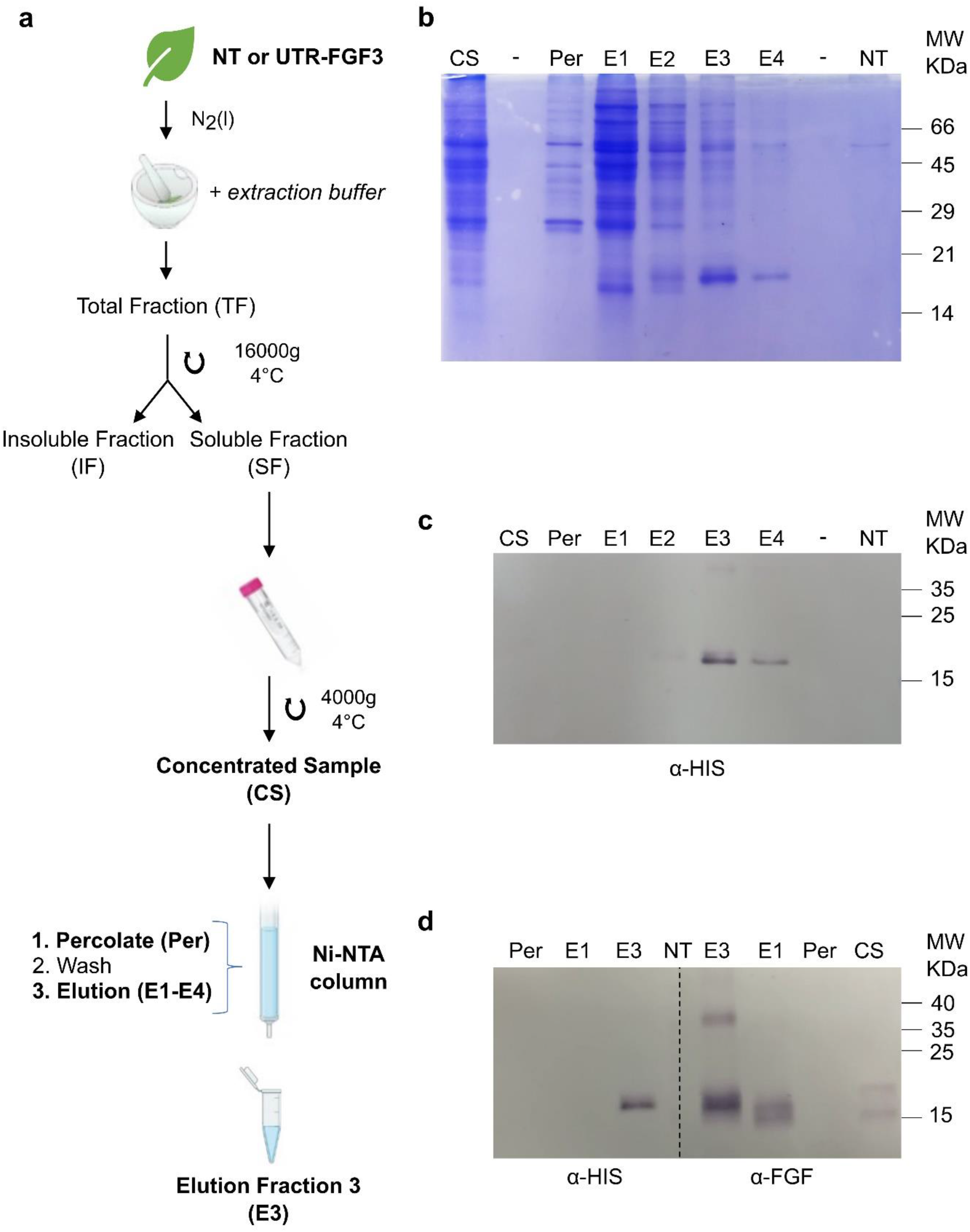
Purification of rhFGFb. **a** Schematic representation of the purification protocol. **b, c** Purification profile. Protein fractions from the purification protocol were analyzed by SDS-PAGE and *coomasie blue* staining (**b**) or *western blotting* with anti-HIS antibody (**c**). **d** Key fractions from the purification protocol were analyzed by western blot with anti-HIS (left) or anti-FGF antibody (right).**CS:** concentrated sample, **Per:** percolate, **E1-E4:** elution fractions, **-:** empty lane, **NT:** E3 elution fraction from NT plant.

Additionally, we analyzed fractions of key purification stages by Western blot comparing anti-HIS and anti-FGF antibodies (Fig. 4d). In the E1 fraction, no band was detected using the anti-HIS antibody. However, a band of lower molecular weight compared to the 6XHIS-tagged rhFGFb was identified using the anti-FGF antibody. This observation suggests that despite adding protease inhibitors during purification, a fraction of rhFGFb undergoes proteolysis, resulting in the loss of the 6xHIS-Tag.

Densitometry analysis revealed that the UTR-FGF3 line produced an estimate of 30 μg rhFGFb per gram of fresh tissue, corresponding to 0.30% of Total Soluble Protein (TSP). Furthermore, a total of 1.3μg rhFGFb per gram of fresh tissue was recovered in the filtered E3 fraction (Suppl. Fig. S3).

### Plant produced rhFGFb is biologically active and induces cell proliferation *in vitro*

To assess the biological activity of the purified recombinant rhFGFb (referred to as plant rhFGFb), we treated HEK293T cells with plant-produced rhFGFb and determined the cell proliferation rate, a routinely assay to determine a biological activity for FGF activity, using the Resazurin staining (Muñoz-Bernart et al. 2023). As a negative control we employed protein extracts derived from NT plants (NT protein extract), while a commercial recombinant hFGFb (Gibco FGF, PHG0026) served as positive control. As expected, treatment with commercial rhFGFb induced the proliferation of HEK293T cells. Notably, stimulation with plant rhFGFb, resulted in a significant increase in HEK293T cells proliferation compared to cells treated with NT extract (Fig. 5). This result indicates that recombinant rhFGFb purified from transplastomic leaves is biologically active and can induce *in vitro* the proliferation of embryonic cells.

**Fig 5.**
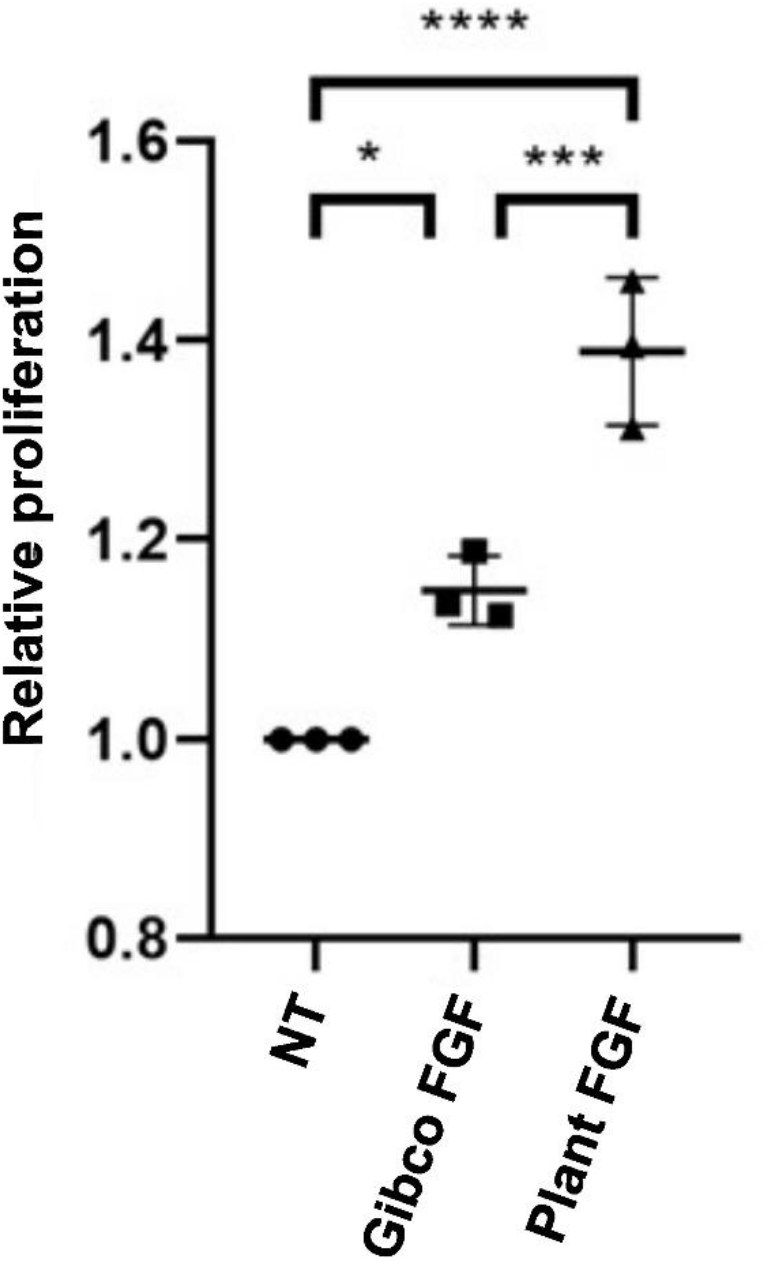
Proliferation assay in HEK293T cells. Rezazurin proliferation assay of HEK293T cells at 24 hs post-induction with non-transformed plant extract (NT), purified plant rhFGFb or commercial FGF (Gibco FGF, PHG0026). Comparison of means of NT control vs rhFGFb (p = 0.0182) or Commercial rhFGFb, PHG0026 (p= 0.0269) by one way ANOVA (n=3).

## Discussion

Growth Factors have gained increasing interest due to their critical roles in regenerative medicine, stem cell research, and more recently cellular agriculture (Venkatesan et al. 2022). One of the most relevant growth factor for stem cell research is the basic Fibroblast Growth Factor (FGFb), which is widely used for culturing human embryonic stem cells (hESCs) (Levenstein et al. 2006).

*E. coli* serves as a commonly employed host for producing recombinant proteins and is the system of choice for commercially produced recombinant hFGFb. However, incorrect protein folding or protein aggregation in inclusion bodies can limit recombinant protein yield in this platform (Bhatwa et al. 2021). Several efforts have been made to optimize the expression of rhFGFb in this system by using peptide fusion partners that increase protein solubility, such as the human protein disulfide isomerase domains (Song et al. 2013), the collagen-like protein (Scl2-M) from *Streptococcus pyogenes* (Rahman et al. 2020), or the Fh8 peptide from *Fasciola hepatica* (Kim et al. 2021). Nevertheless, the risk of endotoxins contamination and the requirement of extensive downstream purification for their removal remain important drawbacks of this system (Wilding et al. 2019).

Plant molecular farming represents an alternative method that enables cost-effective and scalable production while eliminating the risk of contamination with animal pathogens (Buyel 2019). Over the past decade, various approaches and transformation methods have been explored to produce recombinant hFGFb in plants (summarized in Suppl. Table S1). The nuclear stable expression of rhFGFb in the endosperm of seeds has been explored to improve protein stability and streamline downstream purification processes. A rhFGFb with conserved biological activity was expressed in soybean seeds, yielding expression levels of 2.3% TSP (Ding et al. 2006). Furthermore, a biologically active rhFGFb was successfully accumulated in rice endosperm, achieving levels of 185.66 mg/kg of grain. However, the purification of rhFGFb was hindered by insolubility of the recombinant protein and the final recovery was 4.49% of the total rhFGFb expressed in seeds (An et al. 2013). Another optimization strategy involved fusing rhFGFb to oleosin, allowing the expression of a functional rhFGFb within oil bodies from *N. benthamiana* seeds. The rhFGFb expression achieved with this approach was up to 89.95 ng rhFGFb per microliter of oil body, and the oleosin-hrFGFb proved to promote cell proliferation and wound healing (Yang et al. 2018). However, it is noteworthy that the recovered recombinant protein was fused to oleosin and no additional treatments were carried out to obtain a purified rhFGFb. A recent study demonstrated the feasibility of expressing rhFGFb in transplastomic tobacco plants obtaining expression levels of 0.1% TSP (Wang et al. 2016). However, the expression yields obtained by Wang et al. resulted lower than in the afore mentioned works and the biological activity of the recombinant protein was not further investigated.

In our study, we successfully expressed and purified a biologically active rhFGFb from transplastomic tobacco leaves. To assess its functionality, we conducted a proliferation assay using HEK293T cells, a commonly used cell line in biological research that express all of the endogenous Fibroblast Growth Factor Receptor (FGFR) isoforms (Xiao et al. 2016). The accumulation levels of rhFGFb achieved in transplastomic plants UTR-FGF reached the 0.30% TSP, which resulted in a higher yield compared with the work from Wang et al. Although the recombinant protein accumulation levels obtained in our study were sufficient for both rhFGFb purification and the assessment of its biological activity, they did not attain the notably high accumulation levels observed for certain proteins previously expressed by plastid genome transformation (Lentz et al. 2010).

The redox environment of the chloroplast stroma has shown to negatively impact the stability of recombinant proteins, as demonstrated in the case of recombinant hEGF (Wirth et al. 2006). Additionally, a previous study compared the accumulation of a protein with disulfide bonds in the chloroplast stroma versus the thylakoid lumen, revealing higher in thylakoid lumen (Morgenfeld et al. 2014). Given that the rhFGFb contains a disulfide bond in its structure (Müller et al. 2015), redirecting the recombinant protein accumulation to the thylakoid lumen may positively impact on protein accumulation.

It is worth noting that the expression of foreign proteins in the chloroplast can occasionally lead to unintended pleiotropic effects, including chlorosis, growth retardation, delayed flowering, and male infertility. These effects are dependent on the nature of the expressed protein and are not directly correlated with protein expression (Scotti and Cardi 2014). In this specific study, UTR-FGF transgenic lines exhibited a chlorotic phenotype in mature plants, which persisted throughout all stages of plant development. Additionally, these transgenic lines displayed slower growth and delayed flowering compared to their non-transformed counterparts. To the extent of what is revealed by Southern blot analysis, there is no evidence of possible rearrangements. Taken together, these observations suggest that the expression of the transgene may have impact on chloroplast physiology; however, the two independent lines generated in this study remained viable for the expression and purification of a biologically active rhFGFb. Further optimization of this system can be explored to develop a cost-effective method to produce recombinant hFGFb.

The application of immobilized metal affinity chromatography (IMAC) proved highly effective in obtaining rhFGFb in both sufficient quantity and quality for biological activity assays. Following purification, we recovered approximately 5% of the total rhFGFb expressed in transplastomic lines. It is noteworthy that a proportion of the expressed rhFGFb remained in the insoluble fraction, which negatively impacted on the purification yield. An optimization of the purification protocol could involve cell rupture by sonication, modifications on the extraction buffer, heat-treatment or the use of other protein tags that may enhance protein solubility (Buyel et al. 2015). Our results showed that a fraction of rhFGFb underwent processing that leaded to the removal of the 6xHIS tag, which could also negatively impact the purification yield. One notable limitation when using Poly-His tags for plant-based protein expression is the presence of endogenous plant proteins bearing histidine motifs, that may interfere with the detection and purification of the recombinant protein (Coates et al. 2022). Therefore, a possible strategy to improve the purification process could be the replacement of the 6xHIS tag with other protein tags such as the Cysta-tag, designed by Sainsbury (Sainsbury et al. 2016). The Cysta-tag proved to facilitate efficient purification of plant-produced recombinant proteins using the IMAC technique and could an alternative to improve rhFGFb in transplastomic plant.

Finally, our study demonstrates the successful production of a functional rhFGFb using transplastomic plants, affirming the viability of chloroplast transformation as a suitable method for rhFGFb production. To our knowledge, this represents the first report of the production of an active rhFGFb in tobacco chloroplasts.

## Materials and Methods

### Construction of the chloroplast transformation vector

A 6xHIS tagged version of hFGFb was subcloned in the NdeI and XbaI restriction sites in the chloroplast transformation vector pBSW-UTR, (Wirth et al. 2004; Segretin et al. 2012). The rhFGFb transgene was designed by fusing the human FGFb coding sequence (NCBI Reference Sequence:NG_029067.1) to a N-terminal His-Tag followed by the TEV protease recognition signal (Fig. 1). The sequence was flanked with the restriction enzymes <u>NdeI</u> and <u>XbaI</u> for cloning. This sequence was synthesized by GeneScript gene synthesis service (www.genscript.com) after codon optimization for chloroplast expression. The final vector pUTR-hrFGFb was obtained and confirmed by sequencing.

### Tobacco plastid transformation

Plastid transformation was carried out as described by Svab Z. and Maliga P. (Svab and Maliga 1993), using the PDS 174 1000/He biolistic particle delivery system (Bio-Rad Laboratories, Hercules, CA, USA). Fully expanded leaves from *in vitro* cultured *Nicotiana tabacum* plants (L. cv. Petite Havana) were bombarded with 50 μg of 0.6-μm gold particles (Bio-Rad) coated with 10 μg of plasmid DNA using 1,100 psi rupture discs (Bio-Rad). Spectinomycin-resistant lines were selected on RMOP regeneration medium (Svab et al. 1990) containing 500 mgl-^1^ spectinomycin di-hydrochloride. Regenerated shoots were transferred to spectinomycin-containing Murashige and Skoog basal medium (MS) for root and aerial parts development. To test the transplastomic nature of the shoots, a PCR analysis was carried out using primers that anneal to the 16S plastidic gene (Cl_Fw: 5’ GTATCTGGGGAATAAGCATCGG 3’) and the *aadA* gene (Cl_Rv: 5’CGATGACGCCAACTACCTCTG 3’). A 1,450-bp fragment is expected to be amplified in transplastomic plants, which confirms the integration of the transgenes in the plastome. Selected transplastomic lines were subjected to additional rounds of regeneration in RMOP medium with Spectinomycin, to achieve homoplasmy. Plants from the third regeneration round were transferred to soil and grown under greenhouse conditions to obtain seeds. In the greenhouse, natural light was supplemented 16 h per day by sodium lamps providing 100–300 μmol s^-1^ m ^-2^; the temperature was set at 26°C during day and 19°C in the night.

### Southern blot analysis

Total DNA from leaves was extracted using the CTAB method as described previously by Allen et al (Allen et al. 2006). The extracted DNA (3 μg) was fully digested with NcoI restriction enzyme (New England Biolabs, USA), separated in a 0.8% agarose gel and blotted onto a Hybond+ Nylon membrane (Amersham Biosciences). The membrane was crosslinked with UV and hybridized with a trnI/A probe. The trnI/A probe was generated by random priming with Digoxigenin-11-dUTP. Pre-hybridization was carried out at 48°C for 2 hours in DIG Easy Hyb buffer (Roche). The trnI/A probe was denaturalized for 20 minutes at 68°C and used for hybridization overnight at 48°C. The membrane was washed twice for 10 minutes with buffer SSC 2X, 0.1% SDS at room temperature and three times with buffer SSC 0.1X, 0.1%SDS at 68°C. Lastly, the membrane was washed with Malic Acid Buffer. For the detection of labeled DNA bands the membrane was incubated at room temperature with a blocking solution for an hour and subsequently incubated with anti-DIG antibody (1:10000) for 30 minutes. After washing the membrane with washing buffer (Malic Acid Buffer pH 7.5, 0.3% Tween 20), it was incubated with detection buffer (Tris 0.1M pH 9.5, NaCl 0.1M) for 5 minutes. The chemiluminescent substrate CSPD was added, and the membrane was incubated for 5 minutes at 37°C. The GeneGnome imaging system (Syngene) was used for chemiluminescence imaging.

### Protein extraction and analysis

Total proteins were extracted from leaves of transplastomic and wild-type tobacco plants. Approximately 25 μg of leaf tissue was manually grinded in 125 μL of Laemmli buffer 1X (Laemmli 1970) and heated at 98ºC for 10 minutes. The homogenate was centrifuged for 10 minutes at 16,000 g and the supernatant was recovered. For SDS-PAGE, 20 μL of the remaining supernatant was loaded in a 15-17% polyacrylamide gel and electrophoresed for protein separation. After electrophoresis, gels were stained with *Coomassie brilliant blue* dye or transferred onto a nitrocellulose membrane for antibody detection. The nitrocellulose membranes were stained with *Ponceau dye* to assess the quality of protein transfer. His-tagged rhFGFb was detected using a primary anti-HISx6 mouse polyclonal antibody (Roche Molecular Systems, Inc., USA) or anti-FGF polyclonal antibody (Santa Cruz Biotechnology, Inc, USA). As secondary antibody, we used an anti-mouse antibody (Sigma Aldrich, USA) or an anti-rabbit antibody (Cell Signaling Technology, USA), both linked to alkaline phosphatase. Phosphatase activity was determined by a chromogenic reaction using 5-bromo-4 chloro-3 indolyl phosphate and nitroblue tetrazolium (Sigma, St. Louis, MO, USA) as substrates.

### Protein purification with affinity chromatography

Fresh plant tissue (3 g) was grounded in liquid nitrogen, mixed with 6 mL of extraction buffer (PBS 1X pH 7.4, triton 2% w/w, β-MeOH 0.1 M) and centrifugated for 15 minutes at 16000 g at 4°C. The insoluble fraction was resuspended in PBS 1X and the supernatant was loaded in an Amicon Ultra 10kDa filter (Merck, Germany). The supernatant was centrifugated at 4000 g at 4°C for concentration and buffer exchange with 1 mL of equilibrium buffer containing 50 mM phosphate buffer pH 8.0, 300 mM NaCl and 1 mM cOmplete^™^ protease inhibitor cocktail (Roche Molecular Systems, Inc., USA). The obtained sample was loaded in a Ni-NTA affinity chromatography column (Sigma Aldrich, USA). The column was washed with 4 times with 500 μLwashing buffer (50 mM phosphate-buffer pH=8.0, 300 mM NaCl, 20 mM imidazol) and the sample was eluted in 4 steps with 500 μL elution buffer (50 mM phosphate-buffer pH 8.0, 300 mM NaCl, 250 mM imidazol) per elution. The third elution fraction was filtered with a 0.22 μM syringe filter for sample sterilization. The fractions were loaded into a sterilized Amicon Ultra 10 kDa filter (Merck, Germany) to exchange the buffer for PBS 1X 0.1% BSA endotoxin free. Fractions from the different steps of the purification protocol were analyzed by SDS-PAGE and Western blot as described in the previous section.

### Cell culture

HEK 293T (RRID:CVCL_0063) cell line were acquired from the American Type Culture Collection (ATCC) from colleagues, kept frozen at liquid Nitrogen after received and used in culture for a maximum of 4 months. Mycoplasm contamination was evaluated monthly by PCR. Cells were cultured in complete Dulbecco’s Modified Eagle Medium (DMEM) supplemented with 10% fetal bovine serum (FBS), penicillin 100U.ml^-1^ /streptomycin 100 μg.ml^-1^ and L-glutamine 2 mM (Life Technologies, Inc., USA) in 5% CO2 humidified atmosphere at 37 °C.

### Proliferation Assay

For FGF stimulation, 500 cells were seeded onto 96-well plates 24-hour prior to incubation. The cells were incubated with or without 5 ng/ml FGF (GIBCO, PHG0026) or purified rhFGFb at 37 °C in a 7.0% CO2 incubator for 24 hours as indicated. For Resazurin staining 10% FBS DMEM was supplemented with 30 μM Resazurin (#199303, Sigma Aldrich, USA) and incubated. After 2 hours, fluorescence was measured with a Multimode Plate Reader EnSpire® (Perkin-Elmer, MA, USA).

## Acknowledgements

This work was funded by Grant PICT 2014-3700 from Agencia Nacional de Promoción Científica y Tecnológica. We thank Dr. Clara Isabel Marin Briggiler and Dr. Caroline Ana Lamb for providing the anti-FGF2 antibody, Dr. Jorge Muschietti for providing one anti-HIS antibody, Marina Fumagalli for greenhouse assistance and Dr. Verónica Giammaria for technical support. CM, NB and CFV are PhD fellows of Consejo Nacional de Investigaciones Científicas y Técnicas (CONICET, Argentina); FGM was PhD fellow of CONICET; FBA, CPC, SAW and MES are research scientists of CONICET.

## Author contribution statement

MES, SAW and CPC conceived the project. MES directed the project. MES, SAW, CPC, FBA and CM designed the experiments. CM, NB, FGM and CFV conducted the experiments. All the authors analyzed the data. CM and MES wrote the manuscript first draft and all the authors contributed to its revision. All the authors read and approved the manuscript’s final version.

## Data availability

The data that support the findings of this study are available from the corresponding author María Eugenia Segretin upon reasonable request.

## Conflict of interest

The authors declare no conflict of interest.

## Notes

### Competing Interest Statement

The authors have declared no competing interest.

