## Supplementary material (Tables and Figures) for "Production of biologically active human basic Fibroblast Growth Factor (hFGFb) using *Nicotiana tabacum* transplastomic plants"

### Supplementary Tables

| Species | Transformed genome | Subcellular compartment /organ | Expression levels | Purification yield | Biological activity | Reference |
|---|---|---|---|---|---|---|
| Soybean *(Glycine max)* | Nuclear | Seed | 2.3% Total Soluble Protein (TSP) | ND | Cell proliferation activity on Balb/c 3T3 cells | Ding et al. 2006 |
| Rice *(Oryza sativa)* | Nuclear | Seed | 185.66 mg total FGFb /kg rice 17.74 mg soluble FGFb/kg rice | 8.33 mg FGFb/ kg rice (4.49% of total expressed rhFGFb) | Cell proliferation activity on NIH/3T3 cells and wound healing *in vivo.* | An et al. 2013 |
| Tobacco *(Nicotiana tabacum)* | Chloroplast | Chloroplast stroma | 0.1% Total Soluble Protein (TSP) | ND | ND | Wang et al. 2016 |
| *Arabidopsis thaliana* | Nuclear | Seeds (oil bodies) | 89.95 ng FGFb per microlite of oil body | ND | Cell proliferation activity on NIH/3T3 cells and wound healing *in vivo* | Yang et al. 2018 |
| Tobacco *(Nicotiana tabacum)* | Chloroplast | Chloroplast stroma | 0.3% Total Soluble Protein (TSP) | 1.3µg rhFGFb / gr fresh tissue (5% of total expressed rhFGFb) | Cell proliferation activity on HEK293T cells | Present work |

**Table S1: Expression of recombinant FGFb in different plant-based systems.** They all correspond to rhFGFb except from Wang et al. that neither nucleotide sequence nor origin is described in the manuscript.

### Supplementary figure references

**Fig S1. Identification of transplastomic plants by PCR.** DNA extracted from shoots was used as template in a PCR reaction with specific primers. The annealing sites of the primers are shown (down). A previously characterized transplastomic line was used as positive control (+) and a non-transformed tobacco plant (NT) as a negative control.

**Fig S2. Analysis of recombinant protein rhFGFb solubility.** Different fractions of the protein purification protocol were analyzed by western blot with anti-HIS antibody. The RuBisCo large subunit (Rb-L) was used as a loading control (ponceau red dyed). **TF:** total fraction, **IF:** Insoluble fraction, **SF:** Soluble fraction, **NT:** non-transformed plants.

**Fig S3. Quantification of rhFGFb.** Different dilutions of the purification fractions were loaded in parallel with a standard curve of commercial rhFGF2 (Gibco, PHG0026), and analyzed by western blot with specific anti-FGF antibody. The bands intensities were determined using the Image J software. **FE3:** Filtered Elution fraction 3. **SF:** Soluble fraction. **NT:** E3 elution fraction from NT plants.

### Supplementary Figure 1

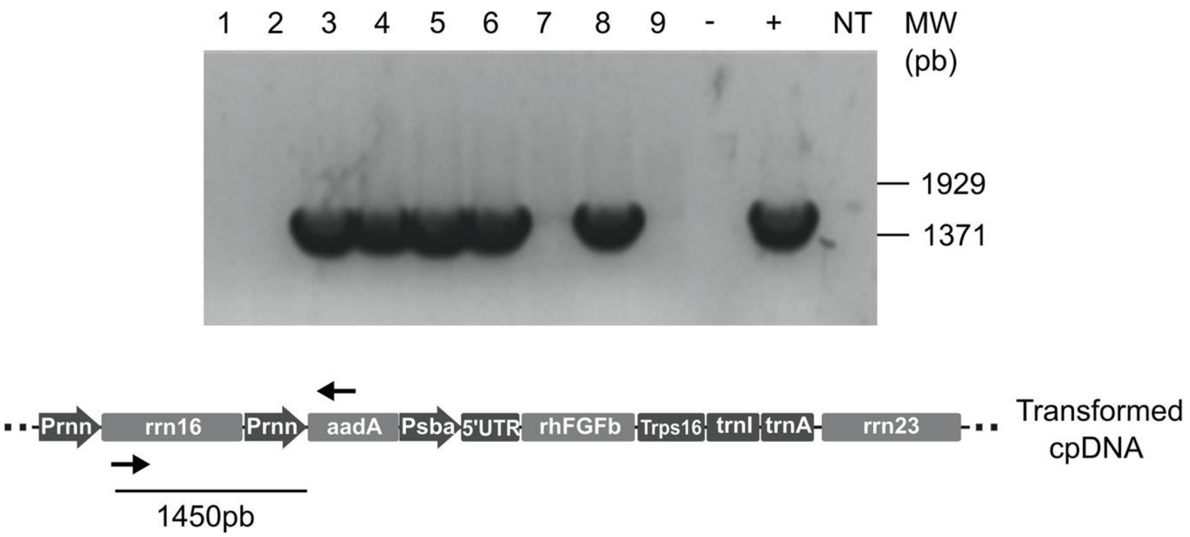

### Supplementary Figure 2

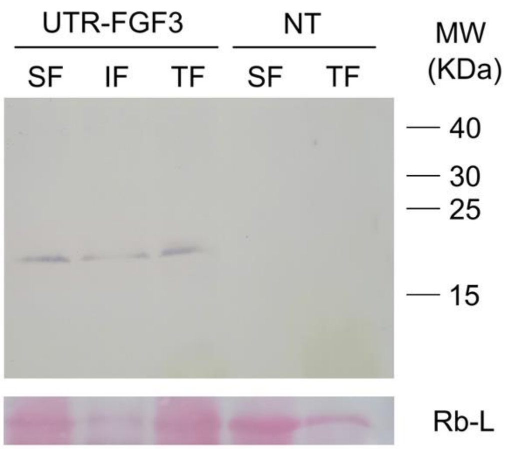

**Supplementary Figure 3**

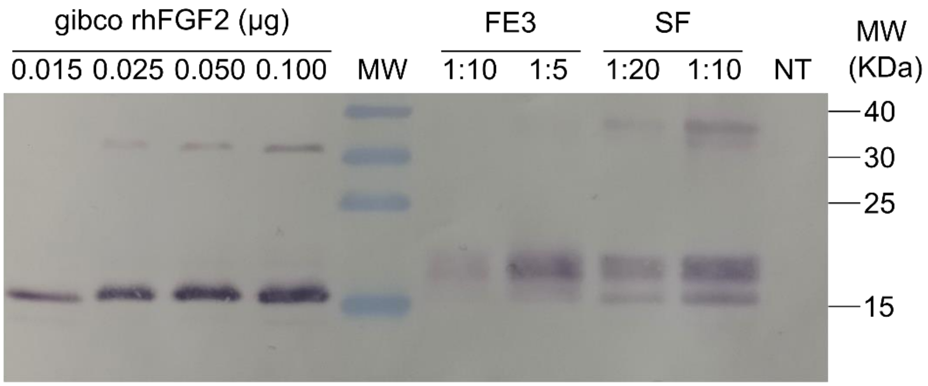
